# DUSP28 is a novel biomarker responsible for aggravating malignancy *via* the autocrine signaling pathways in metastatic pancreatic cancer

**DOI:** 10.1101/313957

**Authors:** Jungwhoi Lee, Jungsul Lee, Chulhee Choi, Jae Hoon Kim

## Abstract

Pancreatic cancer remains one of the most dangerous cancers with a grave prognosis. We previously reported that pancreatic cancer cells can secrete dual specificity phosphatise 28 (DUSP28) to the cultured medium. However, its biological function is poorly understood. Here, we have identified the function of DUSP28 in human metastatic pancreatic cancer. Treatment with recombinant DUSP28 (rDUSP28) significantly increased the migration, invasion, and viability of metastatic pancreatic cancer cells through the activation of CREB, AKT, and ERK1/2 signaling pathways. Furthermore, rDUSP28 acted as an oncogenic reagent through the interaction with integrin α1 in metastatic pancreatic cancer cells. In addition, rDUSP28 induced pro-angiogenic effects in human umbilical vein endothelial cells (HUVECs). Administration of rDUSP28 also produced tumor growth *in vivo.* Notably, sDUSP28 can easily be detected by immunoassay. The results establish the rationale for sDUSP28 as a promising therapeutic target and biomarker for metastatic pancreatic cancer patients.

## Introduction

Pancreatic ductal adenocarcinoma, also known as pancreatic cancer, is one of the most aggressive diseases in the world. The five year survival rate is below 7% and median survival is approximately 6 months. While relatively low in prevalence, pancreatic cancer is the fourth leading cause of cancer-related death [1-3]. More seriously, the prognosis of pancreatic cancer patients remains unchanged, in spite of significant improvements in overall survival rates of other cancers [4]. Effective blockade of pancreatic cancers is urgently required.

One of the major features of pancreatic cancer is the high resistance to conventional cancer therapies including chemotherapy and radiation therapy. Another serious feature of pancreatic cancers is its early distant metastasis and locally abnormal progression, which contributes to the relative rarity of surgery [5, 6]. The more fundamental reason for the extremely poor prognosis may be that only few patients undergo surgical operations on lack of diagnosis [7]. Thus, early diagnosis is of vital significance.

To date, the diagnosis of pancreatic cancer after the onset of symptoms includes the use of computed tomography (CT) scan, endoscopic ultrasound (EUS), and endoscopic cholangiopancreatography (ERCP). Although these techniques are powerful, their ability to diagnose early-stage pancreatic cancer has proven disappointing [8]. The elucidation of genetic abnormalities as a tissue biomarker has also been developed for pancreatic cancer patients. Examples are *KRAS, P16^INK4A^*, *p53*, and *SMAD*. In addition, human serum or plasma markers have been continuously developed due to their easier accessibility for diagnostic testing than tissue samples. In the last two decades, potential biomarkers examined for pancreatic cancer diagnosis include CA19-9, DUPAN-2, CAM17.1, TPS, SPan-1, TAT1, POA, YKL-40, TUM2-PK, and MMP. CA19-9 is the superior marker. Still, it is not the perfect, with variable and sometimes poor sensitivity (41-86%) and specificity (33-100%) [9, 10]. The search continues for a truly effective diagnostic marker for pancreatic cancer patients.

Dual specificity phosphatases (DUSPs) are a heterogeneous group of protein phosphatases that regulate mitogen-activated protein kinases (MAPKs). DUSPs have been implicated as critical modulators of intracellular signaling pathways involved in various diseases including malignant cancers [11, 12]. So far, 25 DUSP genes have been listed in Human Genome Organization databases. These can be subdivided into three groups by subcellular expression. Class I DUSPs including DUSP1, 2, 4, and 5 are localized to the nucleus, while Class II DUSPs including DUSP6, 7, and 16 are found in the cytoplasm. Class III DUSPs including DUSP8, 9, and 10 can be localized in the nucleus or the cytoplasm [13-15]. Interestingly, we previously reported that DUSP28 can be detected in cultured medium of Capan-1 human metastatic pancreatic cancer cells [16]. Although we have proposed the first expression type of DUSPs, its underlying understanding remains poor.

In the present study, we provide the first evidence that metastatic human pancreatic cancers produce the functionally secreted DUSP28 protein, which is critical in metastatic pancreatic cancer malignancy *via* autocrine signaling pathways. Our results provide a rationale for secreted DUSP28 as a therapeutic target and an effective biomarker for metastatic pancreatic cancer patients.

## Results and Discussion

### Effect of DUSP28 treatment in human metastatic pancreatic cancer cells

We have previously reported that Capan-1 human metastatic pancreatic cancer cells specifically secrete DUSP28 (sDUSP28) to the cultured medium compared to six other pancreatic cancer cell types [16]. To confirm the release of DUSP28 in metastatic pancreatic cancer cells, we additionally used CFPAC-1 human metastatic pancreatic cancer cells and SNU-213 cells, which weakly express DUSP28, for sandwich-ELISA and immune-precipitation [17, 18]. DUSP28 were clearly detected in cultured supernatants of Capan-1 and CFPAC-1 cells, but not SNU-213 cells (Supplementary fig. 1A, B). Next, we prepared recombinant DUSP28 (rDUSP28) to evaluate the effects of the DUSP28 in features of human metastatic pancreatic cancer. As shown in figure 1A, rDUSP28 showed a significant and dose-dependent effect on migration of Capan-1 and CFPAC-1 cells. Treatment with maximal concentration of DUSP28 (25 μg/mL) for 6 hours increased Capan-1 and CFPAC-1 cell migration by approximately 1.34- and 1.33-fold compared to control cells; in contrast, denatured DUSP28 (25 μg/mL) had no effect on migration of Capan-1 and CFPAC-1 cell. Exogenously treated with DUSP28 also significantly induced the invasion of Capan-1 and CFPAC-1 cells in a dose-dependent manner; in contrast, denatured DUSP28 had no effect on invasion of Capan-1 and CFPAC-1 cell (Fig. 1B). In addition, the viability of Capan-1 and CFPAC-1 cells was significantly increased in a dose-dependent manner by sDUSP28 treatment; however, denatured rDUSP28 had no effect on viability of Capan-1 and CFPAC-1 cell (Fig. 1C). To understand the mechanism by which sDUSP28 aggravated the malignancy of human metastatic pancreatic cancer cells, we examined the signal transduction pathways using phospho-MAPK array. Exogenous rDUSP28 treatment induced a significant increase in phosphorylation of various MAPK molecules including AKT2 (1.45 fold), CREB (1.25 fold), ERK1 (1.56 fold), ERK2 (1.34 fold), heat shock protein (HSP)27 (1.14 fold), and target of rapamycin (TOR) (1.58 fold) in Capan-1 cells compared to control cells (Fig. 1D). To further analyze the detailed mechanism of DUSP28 treatment in Capan-1 cells, the levels of phosphorylated CREB, AKT, and ERK1/2 were examined with different lengths of time in rDUSP28 treatment. Exogenous DUSP28 treatment increased phosphorylation of CREB, AKT, and ERK1/2 in a time-dependent manner (Fig. 1E). *In silico* protein networks of regulated molecules by exogenously treated with rDUSP28 were constructed with direct target proteins and their interaction partners through the protein-protein interaction data from The Human Protein Reference Database [19]. According to the protein-protein interaction (PPI) network, AKT2, CREB, ERK1, ERK2, HSP27, and TOR regulated by rDUSP28 treatment markedly constructed the anti-cancer signaling pathway including negative regulation of apoptosis and positive regulation of migration (Fig. 1F and supplementary table). These results indicated that secreted DUSP28 is scattered and functionally activates the malignancy in human metastatic pancreatic cancer cells through the activation of MAPK signaling pathways.

**Figure 1.**
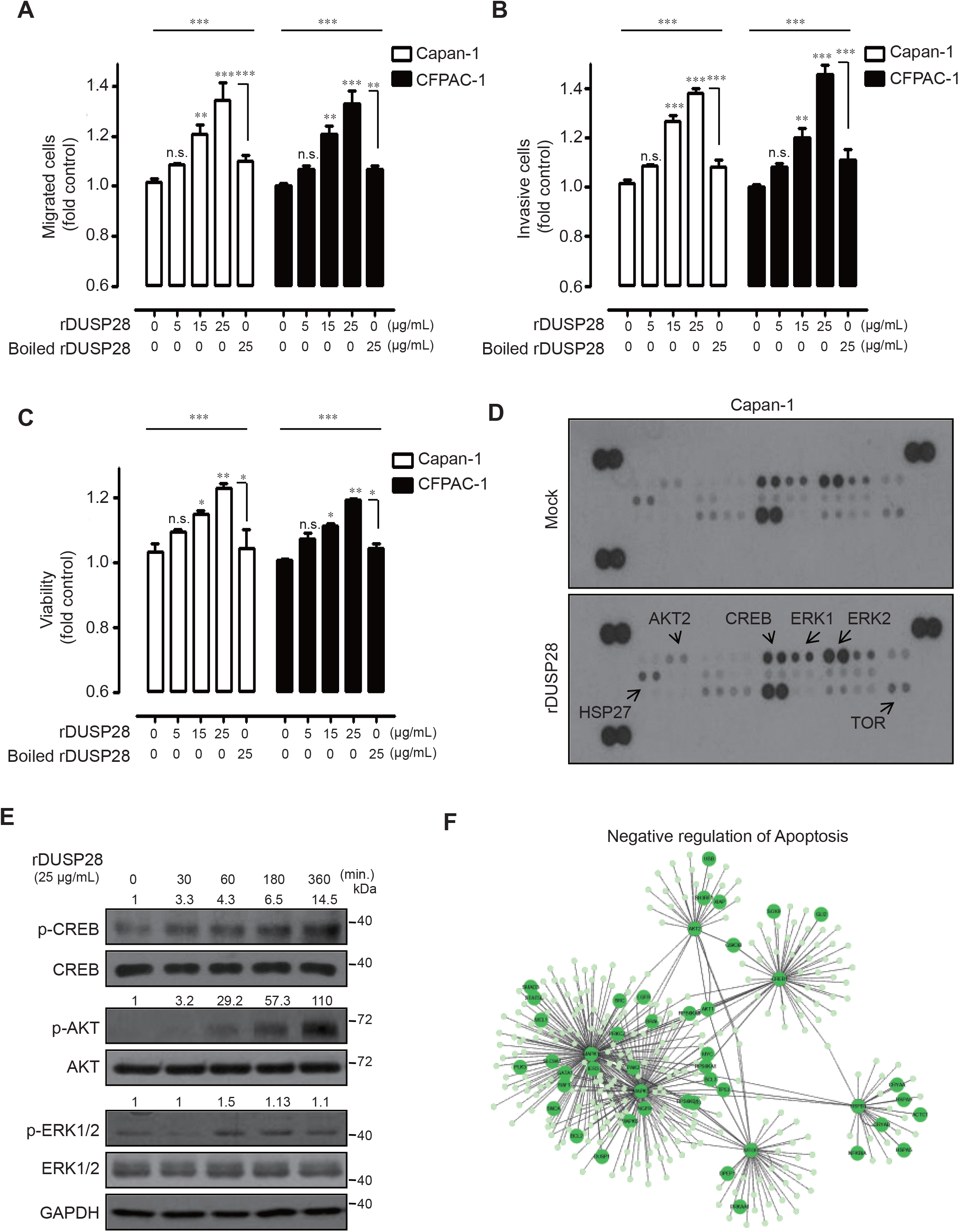
Autocrine signaling mode of DUSP28 in human metastatic pancreatic cancer cells. **A.** Capan-1 and CFPAC-1 cells were incubated with dose-dependent recombinant DUSP28 (rDUSP28) or denatured rDUSP28 for 6 h. Migration was evaluated using the Transwell migration assay (*n* = 3; Tukey’s post-hoc test was applied to detect significant group effects as determined by analysis of variance, *P* < 0.0001; asterisks indicate a significant difference vs. 0% inhibition, *\*\*P* < 0.01, *\*\*P* < 0.01, n.s., non-significant). **B**. Capan-1 and CFPAC-1 cells were incubated with various concentrations of rDUSP28 or denatured rDUSP28 for 24 h. Invasion was evaluated using the Transwell invasion assay (*n* = 3; Tukey’s post-hoc test was applied to detect significant group effects as determined by analysis of variance, *P* < 0.0001; asterisks indicate a significant difference vs. 0% inhibition, *\*\*P* < 0.01, *\*\*P* < 0.01, n.s., non-significant). **C.** Capan-1 and CFPAC-1 cells were incubated with various concentrations of rDUSP28 or denatured rDUSP28 for 72 h under serum-free cultured condition. Viability was measured by the WST-1 assay (*n* = 3; Tukey’s *post hoc* test was applied to significant group effects in ANOVA, p < 0.0001; asterisks indicate a significant difference compared with 0% inhibition, *\*P* < 0.05, *\*\*P* < 0.01, n.s., non-significant). **D.** Capan-1 cells were incubated with rDUSP28 (25 μg/mL) for 24 h. Phospho MAPK array was used to determine differences in rDUSP28-treated, or control human pancreatic cancer cells. Expression changes of various molecules are indicated by the arrow. Right, Relative pixel intensities for p-AKT2, p-CREB, p-ERK1, p-ERK2, p-HSP27, and p-TOR were measured by densitometry analysis using ImageJ analysis software and compared to positive control spots. **E.** Capan-1 cells were incubated with rDUSP28 (25 μg/mL) for various times, and the cell lysates were subjected to Western blot analysis using antibodies specific for phospho-CREB, total CREB, phospho-AKT, total AKT, phospho-ERK1/2, total ERK1, and GAPDH. Relative pixel intensities for p-CREB, p-AKT, p-ERK1/2 were measured using pCREB/CREB, pAKT/AKT, and pERK1/2/ERK1 by densitometry analysis using ImageJ analysis software. **F.** Protein-protein interaction (PPI) networks of AKT2, CREB, ERK1, ERK2, HSP27, and TOR in pancreatic cancer were constructed using The Human Protein Reference Database.

Several DUSPs have been reported as positive regulators of malignant cancer features including apoptosis, proliferation, cell cycle, migration, and chemo-resistance apart from the functions of classical DUSPs. DUSP3 is up-regulated in cervical and prostate cancers, and functions as an inhibitor of apoptosis [27, 28]. DUSP12 is also amplified in a variety of cancers including neuroblastoma, retinoblastoma, intracranial ependynoma, and chronic leukemia, and participates in cell cycle regulation, inhibition of apoptosis, and activation of migration [29]. Selective over-expression of DUSP23 in MCF-7 human breast cancer cells induces cell proliferation, while knock-down of DUSP23 expression decreases cell proliferation [30]. In addition, enhanced expression of DUSP26 promotes colony formation, while si-DUSP26 transfection reduces cell proliferation [31]. In agreement with previous studies, DUSP28 was uniquely expressed in metastatic pancreatic cancer cells as a secreted form, and aggravated pancreatic cancer malignancy through the activation of the autocrine MAPK signaling pathway.

### Molecular partner with DUSP28 in human metastatic pancreatic cancer cells

We have previously investigated that ITGα1 was highly expressed in various human pancreatic cancer cells and DUSP28 affected the functions of critical molecules in human pancreatic cancer [22, 23]. To validate the correlation of ITGα1 as an interaction partner with DUSP28 in human pancreatic cancer cell, we examined the IP analysis following exogenous rDUSP28 treatment in SNU-213 cells that weakly express DUSP28. ITGα1 was co-precipitated with exogenously treated rDUSP28 in SNU-213 cells compared to which of normal mouse IgG antibody (nmIgG) (Fig. 2A). Transfection of si-ITGα1 decreased both mRNA and protein levels of ITGα1 in Capan-1 cells (Fig. 2B Left, Right). Silencing of ITGα1 significantly diminished migration, invasion, and viability in Capan-1 cells (Fig. 2C, D). In particular, transfection of si-ITGα1 inhibited migration, invasion, and viability stimulated by exogenous rDUSP28 treatment in Capan-1 and CFPAC-1 cells, respectively (Fig. 2E, F). Inhibition of ITGα1 expression was also remarkably decreased the phosphorylation of CREB, AKT, and ERK1/2 activated by exogenous rDUSP28 treatment compared to those in scrambled si-RNA transfected cells, respectively (Fig. 2G). According to the GSE57495 data-set, low ITGα1 expression significantly improves median survival by 391 days compared to that of a high level of ITGα1 (Fig. 2H). These results indicated that DUSP28 aggravates malignancy of human metastatic pancreatic cancer *via* interaction with oncogenic ITGα1.

**Figure 2.**
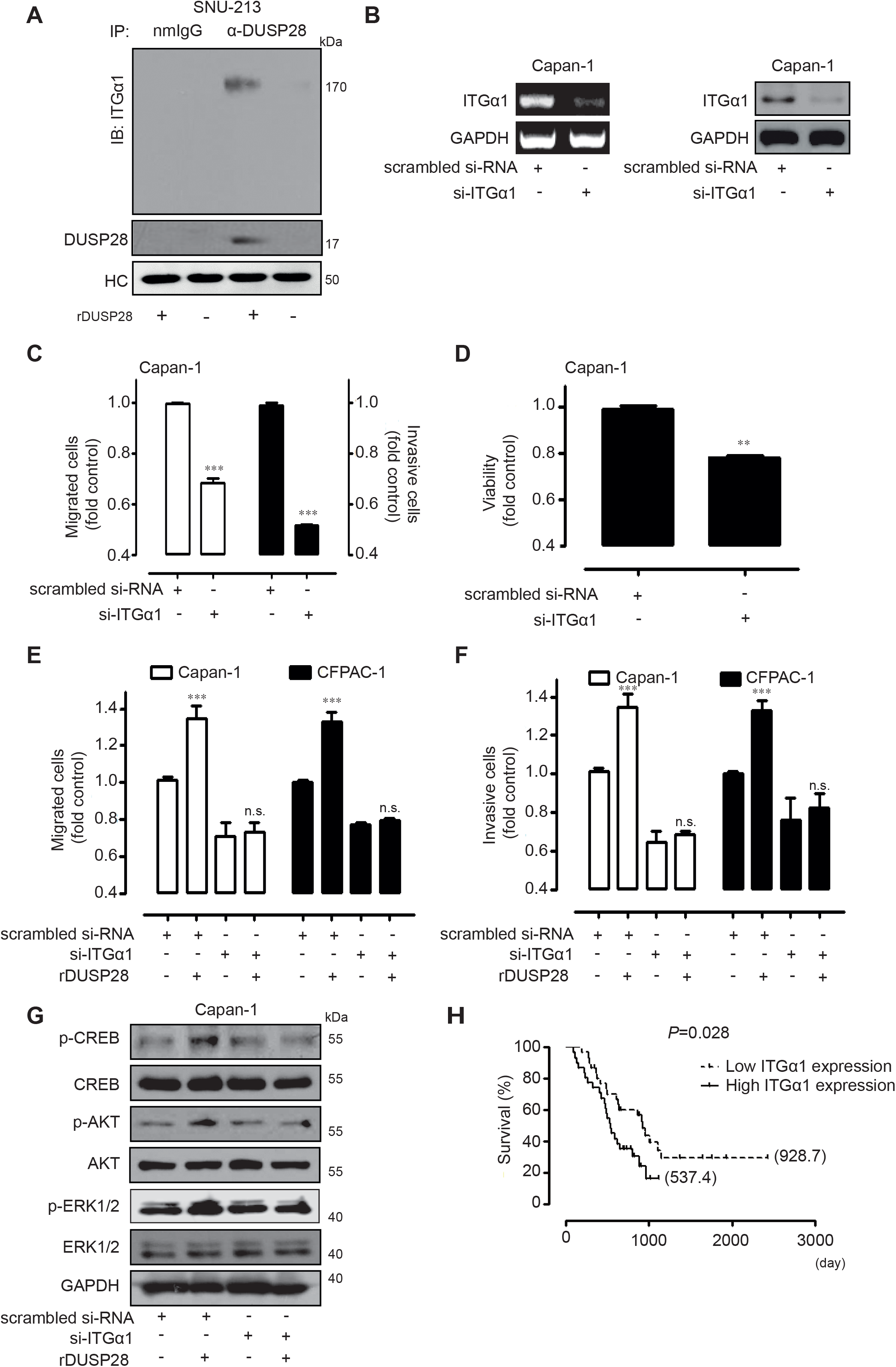
Molecular interaction partner of DUSP28. **A.** ITGα1 and exogenously treated rDUSP28 were precipitated using polyclonal DUSP28 and normal rabbit IgG antibodies in SNU-213 cells. Western blot analysis was performed using monoclonal antibody specific for ITGα1 and DUSP28. Heavy chain (HC) of antibodies was used as a control. **B.** Capan-1 cells were transfected with scrambled or ITGα1-specific siRNA. After 72 h of transfection, *ITGα1* mRNA levels were analyzed by reverse-transcription polymerase chain reaction (RT-PCR, Left) and ITGα1 proteins were subjected to Western blot analysis, Right. GAPDH was used as a control. **C.** Capan-1 cells were transfected with scrambled or ITGα1-siRNAs for 48h. Migrated or invasive cells was evaluated using the Transwell-assay for an additional 6 for migration or 24 h for invasion, respectively (*n* = 3; Tukey’s *post-hoc* test was used to detect significant difference in ANOVA, p < 0.0001; asterisks indicate a significant difference compared with 0% inhibition, *\*\*\*P* < 0.001). **D.** Capan-1 cells were transfected with scrambled or ITGα1-siRNAs for 72h. Viability was measured by the WST-1 assay (*P* value was calculated using the Student’s *t-*test). **E.** Capan-1 and CFPAC-1 cells were transfected with scrambled or ITGα1-siRNAs for 48h. Exogenous rDUSP28 (25 μg/mL) was pretreated for 1h. Migrated cells were evaluated using the Transwell-assay for a 6h (*n* = 3; Tukey’s *post-hoc* test was used to detect significant difference in ANOVA, p < 0.0001; asterisks indicate a significant difference compared with 0% inhibition, *\*\*\*P* < 0.001). **F.** Capan-1 and CFPAC-1 cells were transfected with scrambled or ITGα1-siRNAs for 48h. Exogenous rDUSP28 (25 μg/mL) was pretreated for 1h. Invasive cells were evaluated using the Transwell-assay for a 24h (*n* = 3; Tukey’s *post-hoc* test was used to detect significant difference in ANOVA, p < 0.0001; asterisks indicate a significant difference compared with 0% inhibition, *\*\*\*P* < 0.001). **G.** Capan-1 cells were transfected with scrambled or ITGα1-siRNAs for 48h. Exogenous rDUSP28 (25 μg/mL) was treated for 6h and the cell lysates were subjected to Western blot using antibodies specific for phospho-CREB, total CREB, phospho-AKT, total AKT, phospho-ERK1/2, total ERK1, and GAPDH. **H.** Kaplan-Meier plot of the median survival for pancreatic cancer patients included dependent on differential expressions of ITGα1 (*P* value was calculated using Log-rank (Mantel-Cox) Test, respectively).

Recent work has demonstrated that targeting ITGα1 induced cytotoxicity of pancreatic cancer cells and reduced expansion capacity of collagen *in vitro*, indicating inhibition of ITGα1 sensitizes pancreatic cancer cells to gemcitabine-induced cytotoxicity. In addition, TGF-β cooperated with collagen ECM protein through the ITGα1 dependent mechanisms to establish viable mesenchymal population that are required for systemic spread and survival of pancreatic cancer cells [38]. Another report demonstrated that enhanced DUSP12 induced cellular migration and protection from apoptosis through the upregulation of ITGα1 and the hepatocyte growth factor receptor, c-met. In agreement with previous works, silencing ITGα1 significantly decreased migration, invasion, and viability of human metastatic pancreatic cancer cells. In addition, we revealed that ITGα1 cooperated with exogenous DUSP28 like interaction between growth factor and its receptor to promote metastatic pancreatic cancer malignancy via induction of MAPK signaling pathway such as CREB, AKT, and ERK1/2. Furthermore, we demonstrated that enhanced ITGα1 expression was critical for prognosis of human pancreatic cancer, suggesting that ITGα1 promote the malignancy of pancreatic cancer cooperating with sDUSP28

### Effects of DUSP28 treatment in human umbilical vein endothelial cells (HUVECs)

We next tested whether DUSP28 can induce pro-angiogenic effects in HUVECs due to its external-cellular action potential. As shown in fig. 3A, rDUSP28 treatment significantly induced a HUVECs migration as compared to the control cells in a dose-dependent manner. Treatment with rDUSP28 also significantly increased on the viability of HUVECs as the treated dosages increased (15 μg/mL: 1.12-fold; 25 μg/mL: 1.19-fold) (Fig. 3B). Since endothelial cell morphogenesis is critically required for angiogenesis [20], we next examined the effect of DUSP28 on tube formation in HUVECs using a capillary-like tubular structure that formed on matrigel. Quantitative evaluation of tube formation by counting the junctions of branches revealed that exposure to rDUSP28 significantly increased the number of junctions of the tubular structure compared to control treatment in a dose-dependent manner (15 μg/mL: 3.16-fold; 25 μg/mL: 8-fold) (Fig. 3C). To understand the functional mechanism of the pro-angiogenic effect of rDUSP28 in HUVECs, we examined the time course of rDUSP28-induced signal transduction. We have previously reported that aqueous extraction of *Citrus* unshiu peel induces the pro-angiogenic effect through the activation of FAK and ERK1/2 signaling pathway [21]. Treatment with rDUSP28 (25 μg/mL) time-dependently increased the levels of phosphorylated focal adhesion kinase (FAK^Y397^) and ERK1/2 (Fig. 3D). These results show that treatment with DUSP28 induces pro-angiogenic effects through the activation of the FAK and ERK1/2 signaling pathways.

**Figure 3.**
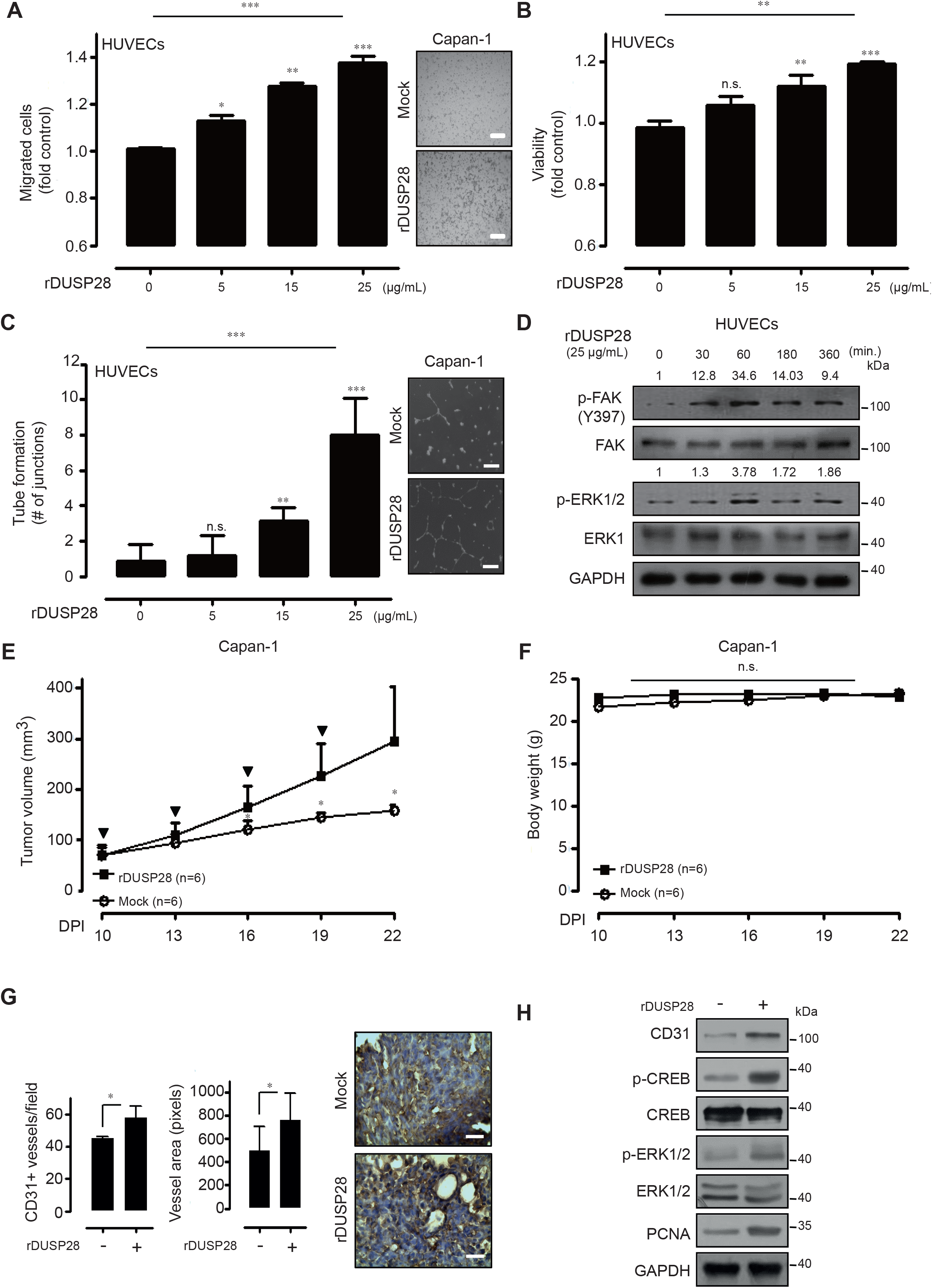
Paracrine signaling mode of DUSP28 in human metastatic pancreatic cancer. **A.** HUVECs were incubated with varying concentrations (0, 5, 15, and 25 μg/mL) of recombinant DUSP28 (rDUSP28). Migration was evaluated using the Transwell migration assay (*n* = 3; Tukey’s post-hoc test was applied to detect significant group effects as determined by analysis of variance, *P* < 0.0001; asterisks indicate a significant difference vs. 0% inhibition, *\*P* < 0.05, *\*\*P* < 0.01, *\*\*\*P* < 0.001, n.s., non-significant, scale bar = 50 μm). **B.** HUVECs were incubated with varying doses of rDUSP28 for 72 h. Viability was measured by the WST-1 assay (*n* = 3; Tukey’s *post hoc* test was applied to significant group effects in ANOVA, *P* < 0.0001; asterisks indicate a significant difference compared with 0% inhibition, *\*\*P* < 0.01, *\*\*\*P* < 0.001, n.s., non-significant). **C.** Capillary like structure (CLS) formation of HUVECs assayed after 16 h of incubation of cells in the varying concentrations of rDUSP28 (data represent the percentage ± SD and are representative of three individual experiments, *\*\*P* < 0.01, *\*\*\*P* < 0.001, n.s., non-significant). **D.** HUVECs were incubated at 25 μg/mL of rDUSP28 in a time-dependent manner, and cell lysates were subjected to Western blot using antibodies for p-FAK and p-ERK1/2. GAPDH was used as loading control. Relative pixel intensities for p-FAK and p-ERK1/2 were measured using pFAK/FAK and pERK1/2/ERK1 by densitometry analysis using ImageJ analysis software. **E.** Effects of rDUSP28 (30 mg/kg) injection in Capan-1 xenograft models (Mock group: *n* = 6 and sDUSP28 injection group: *n* = 6) were measured for 22 days using the formula: V = 0.523 LW^2^ (L = length, W = width). Bold arrows indicate the time of rDUSP28 injection (Tukey’s *post hoc* test was applied to significant group effects in ANOVA, *P* < 0.0001; asterisks indicate a significant difference between the control group and the sDUSP28-injected group, *\*P* < 0.05, DPI indicates the day post injection). **F.** The body weight in each group was measured regularly. **G.** Immunohistochemistry was performed for the number of CD31 positive vessels and vessel areas in control or rDUSP28 treated xenograft tumors. CD31 (brown) was counterstained with hematoxylin (blue) (Magnification: X200, scale bar = 200 μm). **H.** Western blot analysis of control or rDUSP28 treated tumor lysate was probed with anti CD31, p-CREB, p-AKT, p-ERK1/2, and PCNA antibodies. GAPDH was used for loading control.

To verify the autocrine functions of DUSP28 *in vivo*, we prepared xenograft models using Capan-1 cells. Tumor-laden mice were injected intraperitoneally with PBS or rDUSP28 (10 mg/kg) when the tumors reached an average size of approximately 60 mm^3^. PBS-treated Capan-1 xenograft tumors grew to an average size of 157.23 ± 11.33 mm^3^ by 22 days after transplantation, while rDUSP28-treated Capan-1 xenograft tumors grew to an average size of 295.33±108.79 mm^3^ after the same time (Fig. 3E). There was no significant weight loss in either the control or rDUSP28 treated xenograft models (Fig. 3F). As expected, the vessel areas and the number of CD31-positive endothelial cells also were significantly increased in rDUSP28-treated tumors compared to the control group (Fig. 3G). To ascertain the growth stimulating effect of rDUSP28 at the molecular level, we determined the levels of CD31, phospho-CREB, phospho-AKT, phospho-ERK1/2, and proliferating cell nuclear antigen (PCNA) in control and rDUSP28-treated xenograft models. Treatment with 10 mg/kg of rDUSP28 upregulated the levels of phospho-CREB, phospho-AKT, phospho-ERK1/2, CD31, and PCNA compared to the PBS-treated group (Fig. 3H). These results strongly suggest that secreted DUSP28 (sDUSP28) stimulates tumor growth through the activation of the MAPK signaling pathway and angiogenesis *in vivo*.

In addition, to investigate the effect of sDUSP28 inhibition in human metastatic pancreatic cancer cells, we used conditioned medium (CM) containing sDUSP28. We firstly checked the efficiency of si-DUSP28 transfection in Capan-1 cells using RT-PCR and Western blot. Transient transfection of si-DUSP28 strongly reduced the expression of *DUSP28* mRNA in Capan-1 cells (Supplementary fig. 2A, Top) and sDUSP28 in cultured medium (Supplementary fig. 2A, Bottom). Different volumes of si-DUSP28 transfected CM significantly reduced a migration of Capan-1 and CFPAC-1 cells compared to scrambled siRNA transfected CM (Supplementary fig. 2B). Treatment with 10% si-DUSP28 transfected CM (to total volume of cultured medium) also weakly but statistically significantly inhibited the viability of Capan-1 and CFPAC-1 cells (Supplementary fig. 2C). In addition, we examined the effect of si-DUSP28 transfected CM on migration and tube formation of HUVECs using the Transwell migration assay and formation of capillary-like tubular structure that formed on matrigel, respectively. Treatment with different volumes of si-DUSP28 transfected CM significantly showed reduced activities for migration and tube-formation of HUVECs (Supplementary fig. 2D, E). Treatments with different volumes of si-DUSP28 transfected CM also showed weakly, though statistically significant decreases in the viability of HUVECs as the treated dosages increased (Supplementary fig. 2F). These results clearly indicated that blockade of sDUSP28 effectively induces anti-cancer effects in human metastatic pancreatic cancer cells and anti-angiogenesis effects in HUVECs.

Treatment of rDUSP28 also significantly stimulated the angiogenesis *in vitro* and *in vivo*, which indicate that sDUSP28 aggravate the malignancy of metastatic pancreatic cancer *via* dual action mechanisms, autocrine and paracrine signaling pathway, simultaneously. In addition, conditioned medium from si-DUSP28 transfection (less amount of sDUSP28) retarded the basal features of human metastatic pancreatic cancer cells and HUVECs, compared to scrambled siRNA transfected cells. These observations indicate that decent targeting of sDUSP28 might be an advantageous therapeutic strategy for metastatic pancreatic cancer. However, as is the case for most effective therapies, successful targeting of sDUSP28 in metastatic pancreatic cancer will be seriously considered by mining therapeutic reagents, such as therapeutic antibody or small inhibitor. To further understand the effect of sDUSP28 and targeting sDUSP28, various directions and considerable clarifications will be required. Although many DUSPs display enhanced expression and function as oncogenic molecules in various cancers, there are conflicting reports as classical functions of DUSPs [32-35]. Interestingly, different groups have opposing conclusions regarding DUSP26 expression in the same neuroblastoma cell types [36, 37]. Current gaps in knowledge on the controversial functions of DUSPs must be narrow down. Additionally, sDUSP28 signaling has a bi-directional interaction with multiple other pathway or molecules, which include candidate therapeutic targets. Understanding these relationships will significantly stimulate our cogitation to design rational combination therapy.

### Efficacious detection of secreted DUSP28 (sDUSP28)

We explored whether sDUSP28 can be detected in whole blood as a means of diagnosing pancreatic cancers. This exploration was based on the functional recognition of sDUSP28 in cultured cell supernatants. To detect sDUSP28 in whole blood, we prepared xenograft models using Capan-1 (n=4) and SNU-213 (n=4) cells and validated sandwich ELISA using whole blood of Capan-1 and SNU-213 xenograft models. sDUSP28 was specifically detected in whole blood of the Capan-1 xenograft models in a volume-dependent manner, but not in whole blood of the SNU-213 xenograft model and non-tumor bearing mice (Normal) (Fig. 4A). The detection of sDUSP28 in the Capan-1 model’s blood was also significant using Western blot compared to whole blood of the SNU-213 xenograft model and non-tumor bearing mice (Fig. 4B). Furthermore, sDUSP28 could be significantly detected in whole blood from pancreatic cancer patients using sandwich ELISA with a success rate of 83% (5/6) compared to two healthy donors (Supplementary fig. 3). Notably, this easy-to-do technique detected sDUSP28 with an obvious success rate (3/3) in whole blood of metastatic pancreatic cancer patients (C4, C5, and C6) (Fig. 4C). Western blot and IP analysis produced similar results using whole bloods of pancreatic cancer patients and healthy donors (Fig. 4D, E).

**Figure 4.**
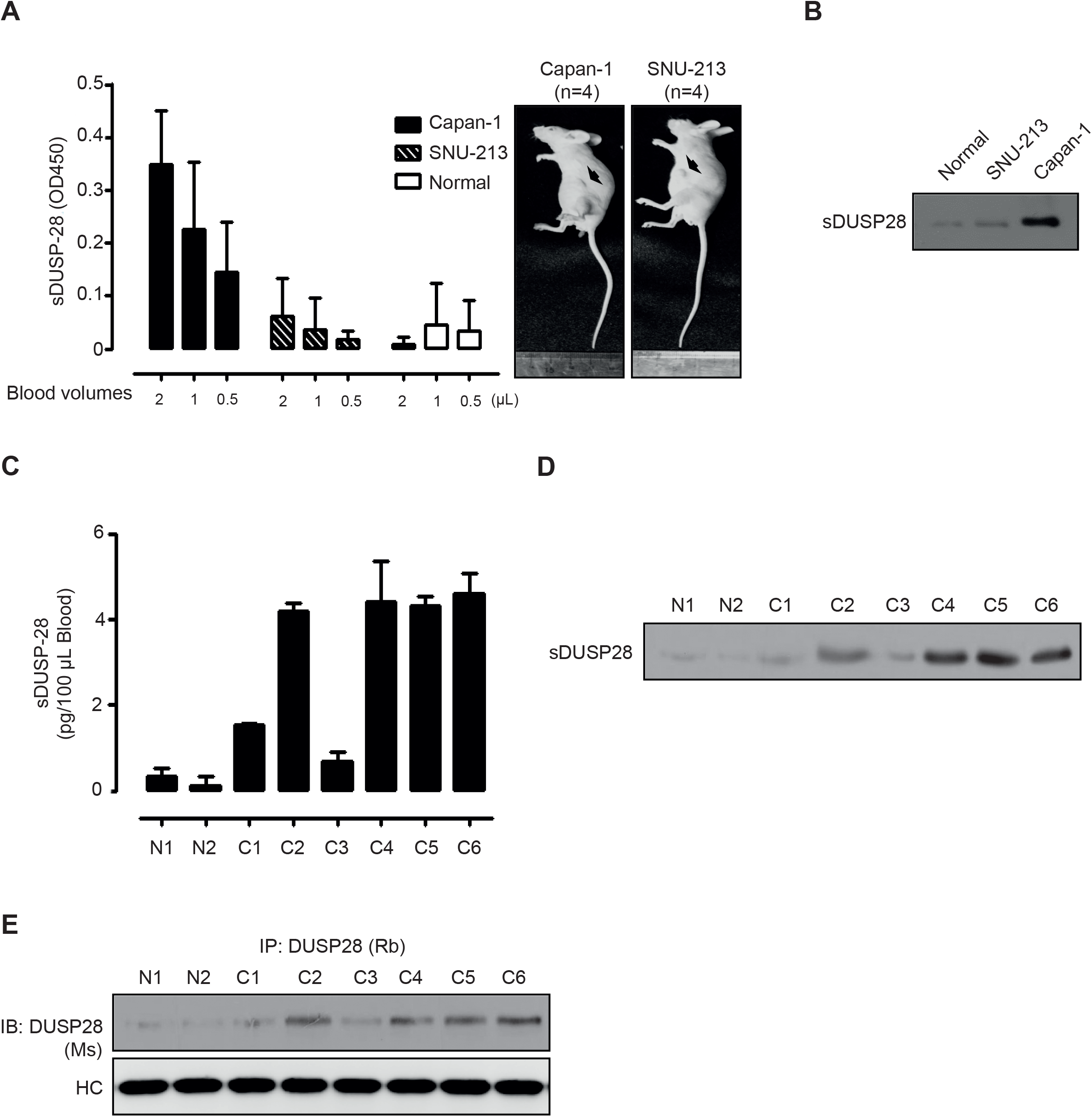
Detection of secreted DUSP28 (sDUSP28) **A.** sDUSP28 was detected by a sandwich ELISA using the varying volumes of whole bloods from Capan-1 and SNU-213 xenograft models (Normal means non-tumor bearing nude mice). **B.** sDUSP28 was detected by Western blot analysis using same volumes of whole blood from Capan-1 and SNU-213 xenograft models (normal, non-tumor bearing nude mice). **C.** Concentrations of sDUSP28 were analyzed by a sandwich ELISA using whole bloods from two health donors (N1 and N2) and six pancreatic cancer patients (C1-C6). **D.** sDUSP28 was detected by Western blot analysis using same volumes of whole bloods from two health donors (N1 and N2) and six pancreatic cancer patients (C1-C6). **E.** sDUSP28 was detected by IP analysis using same volumes of whole bloods from two health donors (N1 and N2) and six pancreatic cancer patients (C1-C6).

Since detection of pancreatic cancer at the earliest stage is the best path to a cure, validation of a truly effective predictive biomarker for diagnosis remains the major challenge for pancreatic cancer. Investigations of existing and new biomarkers to predict prognosis and to stratify therapy are needed. Despite such efforts, the various candidate biomarkers have limited clinical efficiency in diagnosis of pancreatic cancer due to the difficulty in accessing the pancreas, low specificity, and inefficient sensitivity [9]. Currently, CA19-9 is the most promising serum biomarker in pancreatic cancer. However, even CA19-9 is not the perfect biomarker due to its varied sensitivity and specificity [39]. The present data implicate sDUSP28 as a potentially suitable biomarker for metastatic pancreatic cancer. sDUSP28 can be specifically detected in cultured supernatant of metastatic pancreatic cancer cells, Capan-1 and CFPAC-1. We extended the detection to whole blood of metastatic pancreatic cancer cell xenograft mice and pancreatic cancer patients. In particular, sDUSP28 was specifically detected with a significant success rate in whole bloods of metastatic pancreatic cancer patients.

Collectively, these results indicate that sDUSP28 is a critical regulator responsible for metastatic pancreatic cancer malignancy interacting with ITGα1 and its related angiogenesis. Moreover, sDUSP28 might be a potential biomarker for selection of malignant metastatic pancreatic cancer. Thus, the study provides the rationale for development of effective inhibitors against secreted DUSP28 for human metastatic pancreatic cancer patients and secreted DUSP28 might be a promising biomarker for diagnosis of metastatic pancreatic cancer.

## Material and Methods

### Cell culture and reagents

Capan-1 and SNU-213 cells were obtained from the Korean Cell Line Bank (KCLB, Seoul, Korea). CFPAC-1 and human umbilical vein endothelial cells (HUVECs) were purchased from the American Type Culture Collection (ATCC, Manassas, VA, USA). The cells were grown as previously described [16]. Polyclonal antibody for DUSP28 was obtained from Santa Cruz Biotechnology (Santa Cruz, CA, USA) and monoclonal antibody for DUSP28 was purchased from Calbiochem (La Jolla, CA). Antibodies for cAMP response element binding protein (CREB), phospho-CREB (Ser133), protein kinase B (AKT), phospho-AKT (Ser473), extracellular signal-regulated kinase (ERK), phospho-ERK (Thr202/204), proliferating cell nuclear antigen (PCNA), and glyceraldehyde-3-phosphate dehydrogenase (GAPDH) were purchased from Cell Signaling Technology (Beverly, MA, USA).

### Phospho-MAPK array

To demonstrate the intracellular signaling by recombinant DUSP28 treatment, a phospho-MAPK array kit (R&D Systems, Minneapolis, MN) was used according to the manufacturer’s instructions.

### Network analysis and enrichment analysis of chemical targets

Target protein network was constructed with direct target proteins and their interaction partners according to the protein-protein interaction data from The Human Protein Reference Database [19, 40].

### Transfection with small interfering RNA (siRNA)

Transfection of siRNAs was performed using Effectene reagent (Qiagen, Hilden, Germany) as previously reported [21]. Oligonucleotides specific for DUSP28 (sc-94445 and 1120164a/b) were obtained from Santa Cruz Biotechnology and Bioneer (Daejeon, Korea) respectively. Scrambled control (sc-37007) was purchased from Santa Cruz Biotechnology. The efficacy of siRNA transfection was confirmed by Western blot analysis of corresponding proteins.

### Measurement of cell viability

To evaluate cell viability with si-DUSP28 transfection or exogenous treatment with conditioned medium following si-DUSP28 transfection, WST-1 reagent (Nalgene, Rochester, NY, USA) was used as described previously [41]. After a 10-min incubation at room temperature, absorbance was measured at 450 nm using a microplate reader (Bio-Rad, Richmond, CA, USA).

### Trans-well migration assay

Migration assays were performed using a 24-well Trans-well apparatus (Corning, Corning, NY, USA), as described previously [25]. Cells were applied to the upper chamber containing serum-free RPMI and the cells that migrated to the underside of the filter in 6 h were stained. The eluted dye was measured at 560 nm in an ELISA reader (Bio-Rad).

### Tube formation

Tube formation assays were performed as previously described with some modifications [25]. In brief, 250 μl growth factor-reduced matrigel (BD Biosciences, San Jose, CA, USA) was used to coat 24 well plate (SPL Life Sciences, Seoul, Korea) and allowed to polymerize at 37°C for 30 min. HUVECs (3 × 10^4^ cells/well) were suspended in 500 μl serum-free EBM medium containing different dosages of recombinant DUSP28 (rDUSP28). After incubation for 16 h at 37°C, photographs of four representative fields per well were taken using phase contrast microscopy. Endothelial tubes were quantified by counting the number of junctions defined as the origin of two or more branch protrusions.

### Western blot analysis

To evaluate the protein levels of DUSP28 in blood from Capan-1 or SNU-213 xenograft models, Western blot analysis was performed as described previously [25]. Bands were measured by densitometry analysis using ImageJ analysis software.

### RNA preparation and reverse transcript polymerase chain reaction (RT-PCR)

Total RNA was extracted using the TRIzol reagent (Invitrogen, Carlsbad, CA, USA) and converted to cDNA using a RNA PCR kit (Takara Bio Inc., Shiga, Japan) as previously described [16]. RT-PCR analysis was performed using a Takara RT-PCR kit according to the manufacturers’ protocol. Primers for human DUSP28 were obtained from Bioneer (Daejeon, Korea) and GAPDH gene primers were designed using Primer Express 3.0 (Applied Biosystems, Franklin Lakes, NJ, USA). Primer sequences for RT-PCR were as follows: GAPDH (forward primer, 5′-TCACTGGCATGGCCTTCCGTG-3′; reverse primer, 5′-GCCATGAGGTCCACCACCCTG-3′) and DUSP28 (forward primer, 5′-GCGGGATCCATGGGACCGGCAGAAGCTGGGCG-3′; reverse primer, 5′-CCCGCTCGAGTCAAGCCTCAGGGCCCAACCCTAA-3′).

### Enzyme-linked immunosorbent assay (ELISA)

To measure secreted DUSP28 in whole blood of pancreatic cancer patients, blood samples (six from cancer patients and two from healthy individuals) were purchased from Innovative Research (Novi, MI, USA). Ninety six well plates (SPL) were coated with anti-DUSP28 polyclonal antibody (Santa Cruz Biotechnology) at 1 μg/mL. Bound soluble DUSP28 was detected using anti-DUSP28 monoclonal antibody (Abcam, Cambridge, UK) as previously described [42]. The absorbance was measured at 450 nm by using a microplate reader (Bio-Rad).

### Immunoprecipitation (IP)

To detect secreted DUSP28 in whole blood of pancreatic cancer patients, the aforementioned blood samples were incubated with anti-polyclonal DUSP28 antibody, followed by addition of Protein G agarose (Amersham Bioscience, Little Chalfont, UK). Bound proteins were subjected to SDS-PAGE and Western blotting using monoclonal DUSP28 antibody.

### Xenograft tumor model

BALB/c nude mice were obtained from Orient (Seongnam, Korea) at 6-8 weeks of age. Capan-1 (1×10^7^) and SNU-213 cells (1×10^7^) were injected subcutaneously into both sides of the flank as previously described [43]. Once the tumors achieved a size of approximately 300 mm^3^, blood was drawn from xenograft models and non-tumor mice (normal). Body weight was recorded regularly. Animal care and experiments were carried out in accordance with guidelines approved by the animal bioethics committee of Jeju National University (2016- 0049). Animals were anesthetized by chloroform, perfused with PBS, and fixed with formalin (Sigma-Aldrich). Specimens were excised, immersed in formalin and transferred to 30% sucrose (Sigma-Aldrich) solution. Immunohistochemistry was performed using the VECTASTAIN^®^ ABC-kit (Vector Laboratories, Inc., Burlingame, CA, USA) as manufacturer’s recommendation.

### Statistical analyses

All data are presented as mean ± standard deviation. Levels of significance for comparisons between two independent samples were determined by Student’s t-test. Groups were compared by one-way ANOVA with Tukey’s *post hoc* test applied to significant main effects (SPSS 12.0K for Windows; SPSS Inc., Chicago, IL, USA).

## Authors contributions

JW. Lee designed the study, performed the experiments, and drafted the manuscript. J. Lee arranged all data-sets. C. Choi supervised the arrangement of data-sets. J.H. Kim supervised the entire project. All authors discussed data, read the manuscript, made comments and agreed with its contents.

## Acknowledgements

We authors thank Dr. D.G. Jeong (Korea research institute of bioscience & biotechnology, KRIBB) for kindly providing the soluble DUSP28 protein.

## Data Set EV1. DUSP28 protein network

In silico networks of proteins regulated by the treatment with DUSP28 protein were constructed with direct target proteins and their interaction partners through the protein-protein interaction data from The Human Protein Reference Database.

**Supplementary Figure 1. Identification of secreted DUSP28 (sDUSP28) A.**

sDUSP28 was measured by a sandwich ELISA in SNU-213, Capan-1, and CFPAC-1 cells cultured supernatants. **B.** sDUSP28 was precipitated using polyclonal DUSP28 and normal rabbit IgG antibodies in SNU-213, Capan-1, and CFPAC-1 cells cultured supernatants. Bound DUSP28 was subjected to Western blot analysis using monoclonal antibody specific for DUSP28. Heavy chain (HC) of antibodies was used as a control.

**Supplementary Figure 2. *In vitro* effects of secreted DUSP28 (sDUSP28) blockade A.**

Capan-1 cells were transfected with scrambled or DUSP28-specific siRNA. After 72 h of transfection, DUSP28 mRNA levels were analyzed by reverse-transcription polymerase chain reaction (RT-PCR) and sDUSP28 proteins were captured by immunoprecipitation (IP) in cultured medium (C.M.). GAPDH and heavy chain (HC) were used as a control. **B.** Capan-1 and CFPAC-1 cells were incubated with different volumes of scrambled si-RNA C.M. or si-DUSP28 C.M. (0, 5, and 10 % of total volume) for 6h, respectively. Migration was evaluated using the Transwell migration assay (*n* = 3; Tukey’s post-hoc test was applied to detect significant group effects as determined by analysis of variance, *P* < 0.0001; asterisks indicate a significant difference vs. 0% inhibition, *\*P* < 0.05, *\*\*P* < 0.01, *\*\*P* < 0.01, n.s., non-significant, scale bar = 50 μm). **C.** Capan-1 and CFPAC-1 cells were incubated with different volumes of scrambled si-RNA C.M. or si-DUSP28 C.M. (0, 5, and 10% of total volume) for 72 h under serum-free cultured condition, respectively. Viability was measured by the WST-1 assay (*n* = 3; Tukey’s *post hoc* test was applied to significant group effects in ANOVA, p < 0.0001; asterisks indicate a significant difference compared with 0% inhibition, *\*P* < 0.05, *\*\*P* < 0.01, *\*\*\*P* < 0.001, n.s., non-significant). **D.** HUVECs were incubated with different volumes of scrambled si-RNA C.M. or si-DUSP28 C.M. (0, 5, and 10% of total volume) for 6h, respectively. Migration was evaluated using the Transwell migration assay (*n* = 3; Tukey’s post-hoc test was applied to detect significant group effects as determined by analysis of variance, *P* < 0.0001; asterisks indicate a significant difference vs. 0% inhibition, *\*P* < 0.05, *\*\*P* < 0.01, *\*\*P* < 0.01, n.s., non-significant, scale bar = 50 μm). **E.** Capillary like structure (CLS) formation of HUVECs was assayed after 16 h of incubation of cells with different volumes of scrambled si-RNA C.M. or si-DUSP28 C.M. (0, 5, and 10% of total volume), respectively. (data represent the percentage ± SD and are representative of three individual experiments, *\*P* < 0.05, *\*\*P* < 0.01, *\*\*\*P* < 0.001, n.s., non-significant; scale bar = 50 μm). **F.** HUVECs were incubated with different volumes of scrambled si-RNA C.M. or si-DUSP28 C.M. (0, 5, and 10% of total volume) for 72 h. Viability was measured by the WST-1 assay (*n* = 3; Tukey’s *post hoc* test was applied to significant group effects in ANOVA, p < 0.0001; asterisks indicate a significant difference compared with 0% inhibition, *\*P* < 0.05, *\*\*P* < 0.01, n.s., non-significant).

**Supplementary Figure 3. Information of experimental whole bloods**

Clinical information of whole bloods from two health donors (N1 and N2) and six pancreatic cancer patients (C1-C6).

